# Reproducing bench-scale cell growth and productivity

**DOI:** 10.1101/302786

**Authors:** Ariel Hecht, James Filliben, Sarah A. Munro, Marc Salit

## Abstract

Reproducing, exchanging, comparing, and building on each other’s work is foundational to technology advances.^1^ Advancing biotechnology calls for reliable reuse of engineered strains.^2^ Reliable reuse of engineered strains requires reproducible growth and productivity. To demonstrate reproducibility for biotechnology, we identified the experimental factors that have the greatest effect on the growth and productivity of our engineered strains.^3–6^ We present a draft of a Minimum Information Standard for Engineered Organism Experiments (MIEO) based on this method. We evaluated the effect of 22 factors on *Escherichia coli* (*E. coli*) engineered to produce the small molecule lycopene, and 18 factors on *E. coli* engineered to produce red fluorescent protein (RFP). Container geometry and shaking had the greatest effect on product titer and yield. We reproduced our results under two different conditions of reproducibility:^7^ conditions of use (different fractional factorial experiments), and time (48 biological replicates performed on 12 different days over four months).

---

The irreproducibility of experimental results in biotechnology^8^ and bioengineering^9^ must be overcome to realize the potential of biology as a reliable engineering substrate.^1,2^ The synthetic biology community has expressed a desire for experimental protocol standards,^10–12^ supplementing existing standards for genetic modifications.^13^ Minimum information standards have improved reproducibility for qPCR,^14^ microarray,^15^ and genomics^16^ experiments, and a minimum information standard could similarly improve the reproducibility of engineered cell experiments. There have been calls to address reproducibility with reference strains.^10,12^ While reference strains and information standards can and should coexist, information standards are more generalizable, accessible, verifiable, and maintainable.

Biological engineering typically proceeds in three steps: genetically modifying the organism, growing the organism, and assaying its function (**Supplementary Figure S1**). The conditions under which engineered cells are grown can have a large impact on the cell’s performance‐‐the relationship between a genetic modification and its function cannot be fully defined without considering the growth conditions. Here we describe a method to systematically evaluate the effect of experimental factors on growth/productivity of engineered cells, and will recommend the development of a minimum information standard based on this method.

We hypothesize that a sufficient description of experimental factors will enable reproducible performance of engineered cells, and that we can realize this description by (1) building a literature knowledgebase to identify factors, (2) measuring factor effects with an appropriate orthogonal factorial experimental design, and (3) demonstrating that controlling these factors results in reproducible growth and productivity. This description can form the basis of a minimum information standard for growth/ productivity of engineered organisms. We will test our hypothesis with two test cases, *E. coli* engineered to produce the small molecule lycopene, and the heterologous protein RFP.

This paper will focus on experimental factors that define and influence growth conditions for engineered cells. Genetic modifications, both intentional and those arising from evolution, are outside of our scope. Cellular assays are also outside of our scope. We will focus on experimental factors at bench-scale (microtiter plates and shake flasks) in batch culture mode because these formats are often the first step in most bioengineering projects, and unlike larger fermenters, microtiter plates and shake flasks do not allow for continuous monitoring and control of many factors.

The factors that affect cell growth/productivity of engineered *E. coli* can be grouped into three broad categories: (1) media, (2) container, and (3) other factors, including time, environment, selective agents and inoculum (**Supplementary Figure S1**). We identified 32 experimental factors that have been reported to affect cell growth/productivity (**Table 1**).

**Table 1.** experimental factors that have been documented to affect cell growth and productivity. The factors are grouped in 3 broad categories, and can be further broken down to 9 more specific categories. From this list, we selected 22 factors that were practically accessible to us in the laboratory to evaluate. For each factor, we selected two levels, low and high, for evaluation in a fractional factorial design. For quantitative factors, where possible, we added a third center level.

|  | Factor | References | Experimental factor | Low | Center | High |
| --- | --- | --- | --- | --- | --- | --- |
| Media | *Components* |  |  |  |  |  |
|  | Media type | 5,26–33 |  |  |  |  |
|  | Carbon source | 28,31,33–37 | Yeast extract (g/L) | 20 | 24 | 28 |
|  |  |  | Glycerol (g/L) | 3 | 5 | 7 |
|  | Nitrogen source | 33,38 | Tryptone (g/L) | 10 | 12 | 14 |
|  | Inorganic ions | 3,38,39 | Magnesium sulfate (g/L) | 0 | 0.12 | 0.24 |
|  | Manufacturer | 3 | Yeast extract source | Sigma | --- | Millipore |
|  | *Properties* |  |  |  |  |  |
|  | pH | 38,40–44 | pH | 6.7 | 7.2 | 7.5 |
|  |  |  | Buffer capacity (mmol/L) | 70 | 90 | 110 |
|  | Osmolality | 38,44–46 | Osmolality (mmol/kg) | 650 | 750 | 850 |
| Container | *Geometry* |  |  |  |  |  |
|  | Volume | 4,47,48 | Well volume (mL) | 2.5 | --- | 10 |
|  |  |  | Flask volume (mL) | 125 | --- | 250 |
|  | Fill volume | 4,5,32,48–53 | Well fill volume | 10% | --- | 30% |
|  |  |  | Flask fill volume | 7.5% | --- | 15% |
|  | Cover | 52,54–58 | Well cover | Aeraseal | --- | Aluminum |
|  |  |  | Flask cover | Foam | --- | Aluminum |
|  | Well shape | 5,47,50,51 |  |  |  |  |
|  | Well bottom | 5 | Well bottom | Round | --- | Pyramidal |
|  | Flask baffles | 4,6,57–62 | Flask baffles | Unbaffled | --- | Baffled |
|  | Baffle geometry | 4 |  |  |  |  |
|  | Surface coating | 51,63 |  |  |  |  |
|  | Container diameter | 32,41 |  |  |  |  |
|  | *Shaking* |  |  |  |  |  |
|  | Shake speed | 4,5,31,32,47–52,54,63–65 | Shake speed (rpm) | 230 | --- | 460 |
|  | Shake diameter | 32,47,49,51,63,64 |  |  |  |  |
|  | Media viscosity | 4,60,66,67 |  |  |  |  |
| Other | *Time* |  |  |  |  |  |
|  | Time | 5,27 | Time (h) | 24 | 48 | 72 |
|  | *Environment* |  |  |  |  |  |
|  | Temperature | 5,26,27,44,68 | Temperature (°C) | 30 | --- | 37 |
|  | Incubator O<sub>2</sub> concentration | 28 |  |  |  |  |
|  | Lab | 9,37,69–71 |  |  |  |  |
|  | Operator | 71 |  |  |  |  |
|  | *Inoculum* |  |  |  |  |  |
|  | Inoculum concentration | 5,72 | Inoculum amount (cells/mL) | 2 x 10<sup>6</sup> | 8 x 10<sup>6</sup> | 3 x 10<sup>7</sup> |
|  |  |  | Inoculum age (h) | 3 | 16 | 112 |
|  | Glycerol in freezer stock | 73 |  |  |  |  |
|  | Cell phase at preservation | 73 |  |  |  |  |
|  | Time in storage | 74 |  |  |  |  |
|  | *Selective agents* |  |  |  |  |  |
|  | Antibiotic type | 37 |  |  |  |  |
|  | Antibiotic concentration | 26,75 | Antibiotic concentration (µg/mL) | 6.25 | 25 | 100 |
|  | *Inducers* |  |  |  |  |  |
|  | Inducer amount | 5,27,76,77 |  |  |  |  |
|  | Induction time | 5,76 |  |  |  |  |

We evaluated these factors with orthogonal two-level fractional and full factorial experiments. Factorial designs have two main advantages compared with evaluating one factor at a time: increased precision in estimating factor effects with minimal bias from factor interactions, and the ability to detect interactions between multiple factors. Factorial designs also have some limitations: estimates of factors are limited to the levels selected for each factor.^17^

For our first test case, *E. coli* engineered to constitutively produce the small molecule lycopene,^18^ we evaluated the effect of 22 factors on three responses: dry cell mass, titer, and yield (yield is the ratio of titer to dry cell mass, and is a dimensionless, scalable parameter). We chose these 22 factors because they were accessible in our laboratory and relevant to our test strain. For each factor, we selected two levels, low and high; a center level was included when possible (**Table 1**). We quantified the effect of each factor on the responses by the relative effect magnitude, which is the absolute value of the difference between the mean response at the two levels divided by the overall mean response.

Our experimental design consisted of 256 experimental runs organized into three groups. A run is a single combination of experimental growth conditions. Runs were grouped to answer a particular question: which factors have the largest effect on growth/productivity (Group 1), are those effects reproducible under different conditions of use (Group 2), and is a single set of conditions repeatable and reproducible over time (Group 3, see **Methods** for definitions of repeatability and reproducibility). Because it was logistically impossible for us to execute all of the runs in a group in one experiment, groups were divided by factor category (Group 1), factor effect (Group 2) and time (Group 3) for execution in sequential factorial experiments (**Figure 1a** and **Supplementary Data File S1a**).

**Figure 1.**
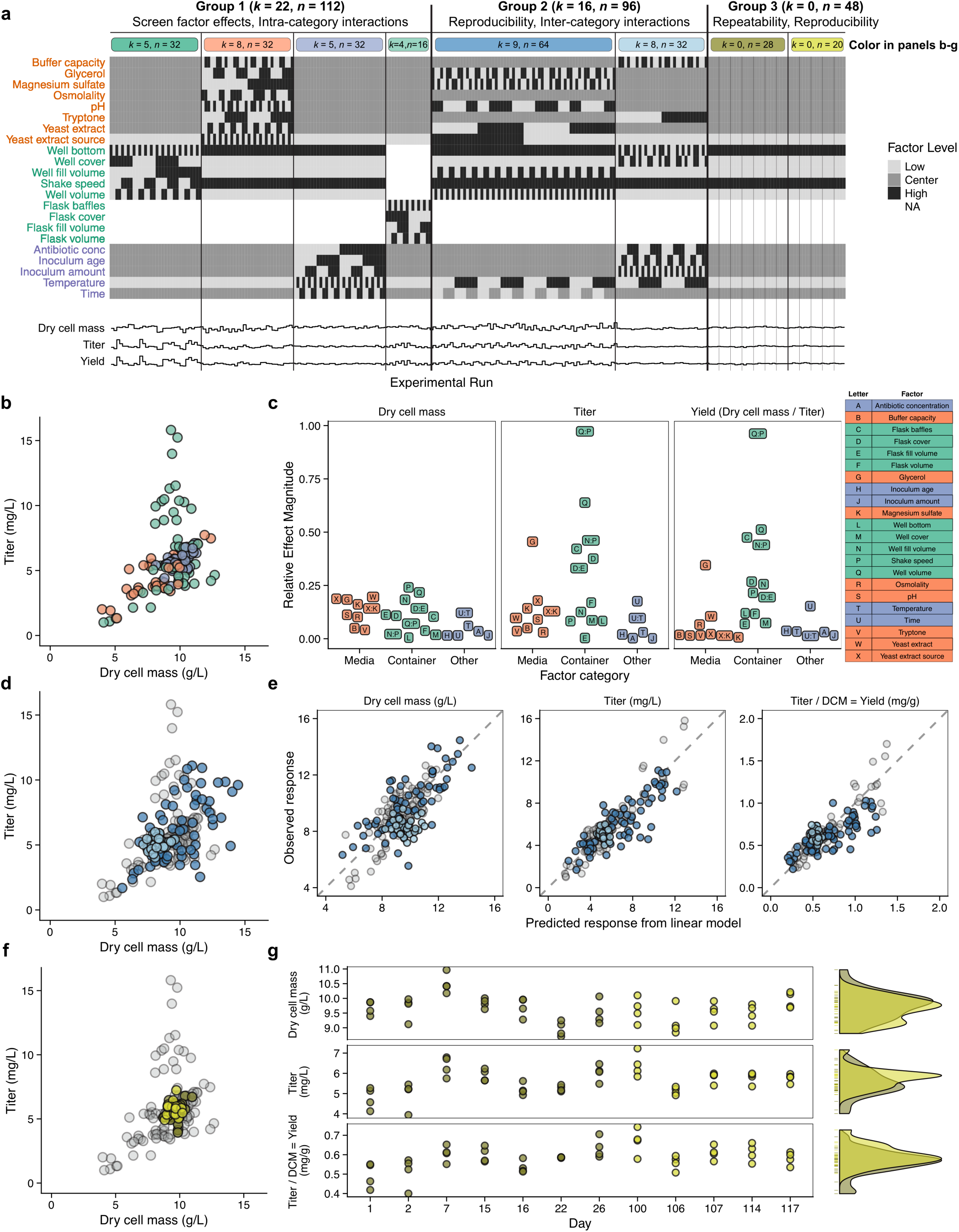
Repeatability and reproducibility of factor effects on cell growth and lycopene productivity. (a) Experimental design table with 22 factor rows, and 256 experimental run columns. Colored bars at the top correspond to color of points in other panels. *k* is the number of factors varied, and *n* is the number of runs, in a group or experiment. The three responses, normalized to range from 0 to 1, are below. (b) Dynamic range of lycopene titer and dry cell mass observed in Group 1 experiments. (c) Relative effect magnitude of all 22 factors on all 3 responses, colored by factor category. (d) Dynamic range of lycopene titer and dry cell mass observed in Group 2 experiments. (e) Factor effects observed in Group 1 are reproducible in Group 2. Grey points are Group 1 data used to train linear model (dry cell mass r^2^ = 0.66, titer r^2^ = 0.85, yield r^2^ = 0.87). Blue points are Group 2 used to test linear model (dry cell mass r^2^ = 0.46, titer r^2^ = 0.71, yield r^2^ = 0.56). (f) Dynamic range of titer and dry cell mass observed in Group 3 centerpoint replicates. (g) Centerpoint replicates plotted as a function of day on which they were run show no trends over time (left). Density plots (smoothed histograms) of response distribution in the first 26 days (dark yellow) and last 17 days (light yellow) overlap (right).

Group 1 screened factor effects and intra-category interactions in 112 runs, divided into four experiments by factor category: media composition, microwell containers, shake flask containers, and other, using 2^8-3^, 2^5^, 2^4^ and 2^5^ orthogonal factorial designs, respectively, with appropriate randomization. These designs allowed us to estimate the main effect of each factor and to detect two-factor interactions within each category. These designs did not allow us to detect interactions between factors in different categories, which we addressed in Group 2.

Varying the factor levels for growing our strain produced a dynamic range of 4-12 g/L for dry cell mass and 1-16 mg/L for lycopene titer (**Figure 1b and Supplementary Figure S2**). We calculated the relative effect magnitude of two-factor interactions for all pairs of factors in each experiment (**Supplementary Data File S1b**), and identified significant two-factor interactions by examining a normal probability plot of the effects (**Supplementary Figure S3**). Particularly interesting two-factor interactions occurred between yeast extract source and magnesium sulfate (supplementing the media with 0.24 g/L magnesium sulfate^3^ eliminated the effect of yeast extract source) and between container factors (non-linear interactions between shaking speed, container volume, and fill volume) (**Supplementary Figures S4-6**).

Container factors and glycerol had the largest effect on strain productivity (**Figure 1c and Supplementary Figure S7**). Container factors primarily affect oxygen transfer into the media, but can also affect exchange of other gases, shear forces, and mixing within the media. The biggest single effect was a the interaction between well volume and shake speed. Titer was more sensitive to container factors than dry cell mass. Dry cell mass and titer were equally sensitive to media factors. Except for glycerol, yield was not sensitive to media factors‐‐changing the media composition affected the total amount of cells that grew in the culture, but not the per-cell productivity. Time and temperature had relatively small effects at the levels used here.

In Group 2, we evaluated the reproducibility of factor effects and screened inter-category interactions in 96 runs. We split the factors into two experiments based on the results from Group 1: 9 factors that had big effects on the responses, and 8 factors that had small effects on the responses, using 2^9-3^ and 2^8-3^ orthogonal fractional factorial designs, respectively (**Figure 1d**). We trained a linear model with the Group 1 data to predict responses from these factors (**Supplementary Methods** and **Data File S1c**). We then tested this model by using it to predict the responses in Group 2 (**Figure 1e**). These results show that the factor effects are predictable by a linear model, reproducible under different conditions of use, and that there were no significant two-factor interactions between factors in different categories (**Supplementary Figure S8** and **Data File S1d**).

In Group 3 we evaluated the repeatability and reproducibility of the growth/productivity at a single set of factor levels (centerpoint levels) in 48 runs. Four replicate runs were performed on each of 12 different days over four months: 7 days in the first month and 5 days in the fourth month (**Figure 1f-g**). The variance within each of the 12 days was homogeneous (**Supplementary Table S1**). We observed that the repeatability within a single day (mean repeatability standard deviation) was 3.5% for dry cell mass, 7.2% for titer, and 8.2% for yield. We observed that the reproducibility between the first and fourth month (reproducibility standard deviation) was 4.9% for dry cell mass, 11.4% for titer, and 11.4% for yield. The distribution of the data from the two months were similar, with month only accounting for 2.3% (dry cell mass), 4.9% (titer) and 8.7% (yield) of the variance with the population, as determined by an analysis of variance (**Figure 1g**). These results show that using our method to identify and control experimental parameters allowed us to reproduce our results over time.

For our second test case, we evaluated the effect of 18 factors on the dry cell mass, titer, and yield of *E. coli* BW25113 engineered to constitutively express the heterologous protein RFP.^19^ We observed similar results as with lycopene, except for different relative effect magnitudes of the container factors on titer (**Supplementary Figures S9-16**). We speculate that these differences may be due to differences in the utilization of oxygen in the biosynthetic pathways of the two products‐‐lycopene is derived from central metabolism, and RFP is a heterologous protein.

In our experiments, we found that the geometry and shaking of the growth container had the largest effect on productivity. These factors warrant special consideration, because they are often tied to large capital expenditures, and can be difficult to change. Given their nonlinear effects, if factor levels cannot be matched in different labs, then expectations about reproducibility should be adjusted accordingly.

We have determined a sufficient description of experimental factors that enabled repeatable and reproducible measurements of growth and productivity of two engineered *E. coli* strains. This demonstrates proof-of-principle of our approach, and is a step towards the creation of a minimum information standard for growth conditions of engineered organisms, which would support interoperability of engineered parts and enable assessment of reproducibility.^10,20^

Experimental growth conditions are frequently specified and documented by free-form text, such as he methods sections of most journals. These unstructured narratives are problematic because it is left to the authors to decide what information to include, and they can be difficult to parse. We propose developing a Minimum Information Standard for Engineered Organism Experiments (MIEO), using a method such as described here, to address this issue. MIEO should be useful for any biological engineer who is planning experiments, reporting results, comparing results, or reproducing results within and between organizations.

To maximize the success of MIEO, we will incorporate lessons about modularity and simplicity learned from previous minimum information checklists. MIEO will be a modular checklist,^16,21,22^ capturing information in nine categories (**Table 2**). The factors that should be included will be different in each experiment (**Supplementary Discussion**). Categories will be designated as required or optional for reporting, similar to other standards.^14,16^ Categories are optional because they can be derived from other categories, or are not applicable in every situation. MIEO is intended to be compatible with any cell type, and any downstream assay, complementing the existing suite of minimum information checklists.^22^

**Table 2.** Minimum Information Standard for Engineered Organism Experiments (MEIO) v0.1, with example for Group 3 single level (centerpoint) replicates (Figure 1a). The nine MEIO factor categories are given, along with whether their reporting is required (R) or optional (O). Factors and their level are reported.

| MEIO Category | Factor | Level |
| --- | --- | --- |
| **Media Components (R)** | Yeast extract | 24 g/L |
|  | Yeast extract source | Sigma Y1625-250G, Lot SLBR9838V |
|  | Glycerol | 5 g/L |
|  | Magnesium sulfate | 0.12 g/L |
|  | Tryptone | 12 g/L |
|  | Potassium phosphate | 2.28 g/L |
|  | Dipotassium phosphate | 12.7 g/L |
|  | Sodium chloride | 6.63 g/L |
| | Water | DI water (18 MΩ-cm) |
| **Media Properties (O)** | pH | 7.2 |
|  | Buffer capacity | 90 mmol/L |
|  | Osmolality | 750 mmol/kg |
| **Container Geometry (R)** | Type | 96 well plate |
|  | Well shape | Square |
|  | Well bottom | Round |
|  | Well volume | 2.5 mL |
|  | Fill volume | 10% (0.25 mL) |
|  | Cover | AeraSeal |
| **Container Shaking (R)** | Shaking speed | 460 rpm |
|  | Shaking diameter | 12.5 mm |
|  | Shaking mode | Orbital |
| **Time (R)** | Growth time | 48 h |
| **Environment (R)** | Temperature | 30 °C |
|  | Relative humidity | 80% |
| **Selective agents (O)** | Antibiotic type | Chloramphenicol |
| | Antibiotic concentration | 25 μg/mL |
| **Inoculum (R)** | Concentration at inoculation | OD<sub>600</sub> = 0.01 |
|  | Age of inoculum at inoculation | 16 hours |
| **Inducers (O)** | (none) | (none) |

One of the key factors that determine the adoption of a minimum information standard is simplicity.^23^ We will aim for simplicity by limiting the standard to nine categories of information, and by including both human-readable/writable (e.g., a table created in word-processing or spreadsheet software) and machine-readable/writable (e.g., XML) implementations. While a machine-readable/writable format has obvious appeal, the cost of adopting such a standard can be prohibitive for some. We have created an example checklist based on the centerpoint experimental conditions used in this paper (**Table 2**).

Standards development is best as a community-driven bottom-up effort, not a top-down prescription.^24,25^ We encourage other members of the biotechnology community to contribute to the development of this standard (http://jimb.stanford.edu/mieo/).

The challenges facing the reproducibility of experimental data in biology are significant. The results shown in this paper demonstrate a method for reproducing key experimental results over time and under different conditions of use. A well-implemented and widely-adopted minimum information standard would improve the repeatability and reproducibility of engineered organism experiments. Experimental reproducibility would advance biological engineering towards becoming a more reliable and predictable engineering discipline.

## Acknowledgments

The authors would like to thank Drew Endy for hosting us in his laboratory and providing helpful insights and conversations, Steve Lund for helpful discussions regarding experimental design, Amar Ghodasara for helpful discussions regarding lycopene assays, Noah Spies for helpful discussions regarding data analysis, and Tobias Meyer and Arnold Hayer for assistance with osmolality measurements. The authors would like to thank the participants and leaders of the Synthetic Biology Standards Consortium for providing inspiration for this work. Certain commercial equipment, instruments, or materials are identified in this report to specify adequately the experimental procedure. Such identification does not imply recommendation or endorsement by the National Institute of Standards and Technology, nor does it imply that the materials or equipment identified are necessarily the best available for the purpose.

## Author Contributions

A.H. developed the concept, designed experiments, collected the data, analyzed the data, and wrote the manuscript. J.F. assisted with designing the experiments and analyzing the data. S.A.M. supervised the work, and assisted with developing the concept and designing experiments. M.S. supervised the work, and assisted with developing the concept, designing experiments, and analyzing the data. J.F., S.A.M., and M.S. edited the manuscript.

## Methods

### Strain engineering

The parent strain for both test strains used in this paper was *Escherichia coli* BW25113 (*Δ(araD*-*araB)567 ΔlacZ4787(::rrnB*-*3) λ*^−^*rph*-*1 Δ(rhaD*-*rhaB)568 hsdR514)* obtained from the Yale Coli Genetic Stock Center (New Haven, CT). For production of lycopene, the parent strain was transformed with plasmid pAC-LYC,^18^ a gift from Francis X. Cunningham Jr (Addgene plasmid #53270). For production of RFP (mRFP1), the parent strain was transformed with plasmid pFAB3992,^19^ a gift from Drew Endy (Addgene plasmid #47823). Plasmid sequences (**Supplementary Data File S3**) and maps (**Supplementary Figure S17**) are available online. All reagents used in this paper with manufacturer, product and lot numbers are available online (**Supplementary Table S2**). Cloning methods are available online (**Supplementary Methods**).

### Cell culture

LB agar plates with the appropriate antibiotic (25 μg/mL chloramphenicol or 50 μg/mL kanamycin) were streaked with a sterile pipette tip from glycerol stocks, and incubated overnight at 37 °C. Plates were stored, wrapped in Parafilm M (Bemis NA, Neenah WI), at 4 °C for up to two weeks, after which they were discarded. All cultures were grown in variants of Terrific Broth (TB), which has a baseline composition of 24 g/L yeast extract, 12 g/L tryptone, 5 g/L glycerol, 0.17 mol/L KH_2_PO_4_ and 0.72 mol/L K_2_HPO_4_.^78^ These values were used as centerpoint values for media composition. We modified this recipe by adding magnesium sulfate to supplement deficient magnesium content in the yeast extract,^3^ and adding sodium chloride to adjust the osmolality.^46^

Every experiment began with a liquid starter culture. A single colony was picked from the agar plate, inoculated into 4 mL of TB with appropriate antibiotic in a plastic-capped 16 × 100 mm glass culture tube (VWR 47729-576), and grown for 16 hours at 37 °C, shaking at 250 rpm with a 25 mm shaking diameter. This culture was diluted with phosphate buffered saline (PBS) to an OD_600_ = 0.5 (as measured in a BRAND semi-micro polystyrene cuvette on a WPA Biowave CO8000 Cell Density Meter), and was then used as the starter culture for inoculating the experimental runs. A fresh starter culture was prepared on each day on which an experimental run was started. All experimental runs started on the same day were inoculated from the same starter culture, except for cultures with different inoculum age. For the inoculum age of 3 hours, a 50 μL aliquot of the 16 hour starter culture was taken and used to inoculate 4 mL of fresh media in a glass culture tube, and returned to the incubator for 3 hours, then removed and diluted to OD_600_ = 0.5. This culture was in exponential phase at the time that it was removed from the incubator. For the inoculum age of 112 hours, after the usual 16 hour incubation, the starter culture was stored at 4 °C for 96 hours, and then removed and diluted to OD_600_ = 0.5 for use.

For cultures with varying pH or varying buffer capacity, media composition was determined by simultaneously solving the Henderson-Hasselbach equation:

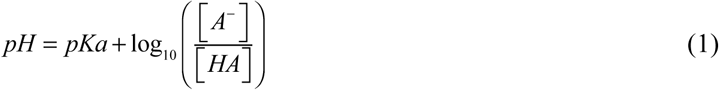

and the equation for buffer capacity:

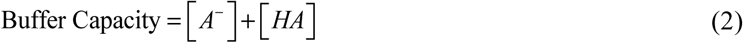

for *[A*^−^] and *[HA]*, where *A*^−^ is the conjugate base, *HA* is the conjugate acid, and *pKa* for phosphate buffers is 6.86.^79^ Solving these two equations for the baseline TB composition gives pH = 7.5 and buffer capacity = 0.89 mM. Adding the remaining media components lowered the pH by 0.3 - 0.5 pH units. In designing the experiments, we aimed for pH of 7.0, 7.5, or 8.0. We measured the pH of the media with a pH meter, and found that the actual pH was approximately 6.7, 7.2 or 7.5, respectively.

For cultures with varying osmolality, the osmolality of all the media components was calculated assuming complete dissociation of ionic species, and the empirically determined osmolality of yeast extract and tryptone of 6 mmol/kg per g/L. Sodium chloride was added to increase the osmolality to the desired level.^46^ The osmolality of 12 different media solutions, as measured by the Wescor Vapro 5520 Vapor Pressure Osmometer, was within 7% of the target value (**Supplementary Table S4**).

The growth of our test strains for the factorial experiments (**Figure 1a and Supplementary Figure 9a**) was performed under the conditions listed in **Table 1** and **Supplementary Data File S1-S2**, and the growth of our test strains for the centerpoint replicates was performed under the conditions listed in **Table 2**. The full experimental design table, with all settings for each run, is available online (**Supplementary Data File S1-S2**). Stocks of each individual media component were prepared at 10x concentration of the centerpoint value with DI water except for magnesium sulfate and dipotassium phosphate, which were purchased. Stocks were sterilized by autoclave except for glycerol, which was sterilized by a 0.22 μm syringe filter (Whatman Puradisc FP30). The position of each run on a plate, and the order in which the runs were set up and sampled, was randomized.

Four different types of container covers were used (**Supplementary Table S2 and Data File S1-S2**). Microwell plates were covered with either the gas-permeable, adhesive AeraSeal membrane (E&K Scientific), or the AeraSeal membrane and a gas-impermeable aluminum foil tape (Bio-Rad). The AeraSeal membrane was first applied to prevent direct contact between the liquid media and the aluminum foil tape. Shake flasks were covered with either the gas-permeable silicone foam covers (Bellco Glass), or two layers of gas-impermeable aluminum foil secured with ParaFilm wrapped around the base of the neck of the flask.

### Assay

Details about assay calibration are available online (**Supplementary Methods**). At the conclusion of the experimental growth time, a 250 μL aliquot from each culture was assayed for dry cell mass and titer. The 250 μL aliquot (in a deep-well 96-well plate) was diluted with 1 mL PBS. A 40 μL aliquot of the diluted cells were transferred into 160 μL of PBS in a clear-bottom polystyrene 96-well plate and mixed.

Absorbance at 700 nm was measured on a Molecular Devices SpectraMax i3 plate reader. For cells expressing RFP, fluorescence was also measured on the plate reader with excitation at 585 nm and emission at 625 nm.

For cells expressing lycopene, the remaining 1210 μL of the diluted cells were transferred to a 1.5 mL microcentrifuge tube, and centrifuged on an Eppendorf MiniSpin microcentrifuge with F-45-12-11 rotor at 1100 *xg* (4000 rpm) for 4 minutes. The supernatant was aspirated, and then lycopene was extracted using a modified version of a previously published protocol.^80^ 250 μL of methanol was added, and the tubes was vortexed vigorously for 5 seconds to break up the cell pellet. A pipette tip was then used to further break up the pellet. 250 μL of acetone was added, and the tubes were vortexed vigorously for 5 seconds again. Then, 250 μL of dichloromethane was added, the tubes were vortexed vigorously for 5 seconds, and then allowed to incubate at room temperature for 10 minutes to complete the extraction of lycopene from the cell pellets. The tubes were centrifuged at 12,000 xg (13,400 rpm) for 5 minutes to pellet the cell debris, and then 200 μL aliquots were transferred to randomly assigned wells on a clear-bottom polypropylene 96 well plate, along with four aliquots of solvent blanks. During the aliquoting process, the plate was covered with a polypropylene plate cover to minimize evaporation, which is a concern when working with small volumes of volatile solvents. Absorbance at 475 nm, 507 nm and 600 nm was measured in the plate reader. Lycopene does not absorb at 600 nm. Absorbance at 600 nm was used to detect the presence of contamination in the samples. The calibration curves were applied to measurements of absorbance at 475 nm and 507 nm to estimate lycopene concentration at these two absorbance peaks, and then averaged to determine the lycopene concentration in the sample.

## Data Analysis

All raw data is available online (**Supplementary Data File S4**). Absorbance and fluorescence measurements were collected through the Molecular Devices SoftMax Pro 6.4 software. All data analysis was performed in R with the FrF2^81^ package for fractional factorial design and analysis, tidyverse^82^ packages for data transformation, and the ggplot2^83^ package for figure generation. All R scripts used to generate the experimental designs and analyze the experimental data are available online (**Supplementary Data File S5**) and at https://github.com/arielhecht/cell-metrics.

Repeatability is the closeness of the agreement between the results of successive measurements of the same measurand carried out under the same conditions of measurement.^7^ Reproducibility is the closeness of the agreement between the results of measurements of the same measurand carried out under changed conditions of measurements (which may include principle of measurement, method of measurement, observer, measuring instrument, reference standard, location, conditions of use, or time).^7^ We evaluated reproducibility under changed conditions of use and time.

In this paper, we define titer as the amount of product produced (mg of lycopene or arbitrary fluorescence units of RFP) per volume of culture. We define dry cell mass as the mass of dry cells per volume of culture. We define yield as the ratio of amount of product produced per mass of dry cells, which is the quotient of titer over yield.

Relative effect magnitude is the absolute value of the difference between the mean response at each level divided by the overall mean response. For one factor, *X_i_*, at two levels, - and +, with one response *Y*, the relative effect magnitude is:

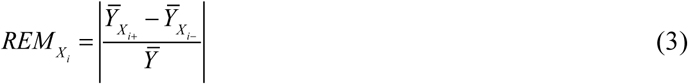

For two factors, *X_i_* and *X_j_*, the relative effect magnitude is:

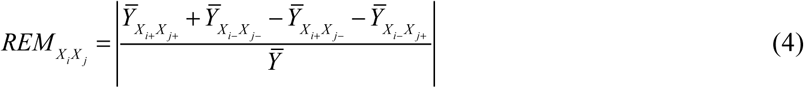

## Data availability

Source data for Fig 1 and Supplementary Figs 2-16 have been provided in Supplementary Data File S4. All other data supporting the findings of this study are available from the corresponding author on request.

